# Co-regulation of transcription by BRG1 and BRM, two mutually exclusive SWI/SNF ATPase subunits

**DOI:** 10.1101/178848

**Authors:** Jesse R. Raab, John S. Runge, Camarie C. Spear, Terry Magnuson

## Abstract

**Background:** SWI/SNF is a large heterogenous multi-subunit chromatin remodeling complex. It consists of multiple sets of mutually exclusive components. Understanding how loss of one sibling of a mutually exclusive pair affects the occupancy and function of the remaining complex is needed to understand how mutations in a particular subunit might affect tumor formation. Recently, we showed that the members of the ARID family of SWI/SNF subunits (ARID1A, ARID1B, and ARID2) had complex transcriptional relationships including both antagonism and cooperativity. However, it remains unknown how loss of the catalytic subunit(s) affects the binding and genome-wide occupancy of the remainder complex and how changes in occupancy affect transcriptional output.

**Results:** We addressed this gap by depleting BRG1 and BRM, the two ATPase subunits in SWI/SNF, and characterizing the changes to chromatin occupancy of the remaining subunit and related this to transcription changes induced by loss of the ATPase subunits. We show that depletion of one subunit frequently leads to loss of the remaining subunit. This could cause either positive or negative changes in gene expression. At a subset of sites the sibling subunit is either retained or gained. Additionally, we show genome-wide that BRG1 and BRM have both cooperative and antagonistic interactions with respect to transcription. Importantly, at genes where BRG1 and BRM antagonise one another we observe a nearly complete rescue of gene expression changes in the combined BRG/BRM double knockdown.

**Conclusion:** This series of experiments demonstrate that mutually exclusive SWI/SNF complexes have heterogenous functional relationships and highlights the importance of considering the role of the remaining SWI/SNF complexes following loss or depletion of a single subunit.

## Introduction

The SWI/SNF complex contains 12-15 subunits [1–3], which are combinatorially assembled to create as many as several thousand biochemically distinct complexes. In development, changes to composition of SWI/SNF can drive developmental progression [4,5]. In cancer, SWI/SNF is among the most commonly found mutations, with mutations observed in as many as 20% of tumors[6,7]. However, these mutations are not equally spread across the subunits. Mutations in the ARID family members (ARID1A, ARID1B, and ARID2) and the ATPase subunits (BRG1 and BRM) are more prevalent than in the other subunits [6,7]. Additionally, in some cancers multiple subunits are mutated. This is the case in hepatocellular carcinoma where mutations have been identified in all three ARID subunits [8–10]. In other cancers, mutations are highly specific, such as BRG1 mutations in Small Cell Carcinoma of the Ovary Hypercalcemic Type (SCCOHT) [11] and SNF5 mutations in malignant rhabdoid tumors [12], and ARID1A in ovarian clear cell carcinoma [13–15]. The varied relationship between mutations in a particular SWI/SNF subunit and the generation of tumors represents a challenge for determining how the SWI/SNF complex affects transcription and other cellular processes during oncogenesis.

We previously used the three ARID subunits as a model to understand how depletion of a single ARID family member affected transcription in HepG2, a well studied liver cancer line [16]. We showed that the ARID family members had a complex relationship that included both cooperative and antagonistic control of genes. Surprisingly, ARID1B and ARID2 had a highly cooperative effect on gene expression, while ARID1A showed both cooperative and antagonistic effects. Additionally, we showed that all three ARIDs bound a highly overlapping set of regions, but that differences in the co-factors at subsets of regions might reflect functionally different complexes. However, in that study we did not determine how loss of one of these subunits affected the genomic occupancy of the remaining subunits. It has been shown that SWI/SNF targeting can be affected by a variety of other chromatin regulating families [17,18], including direct competition with PRC1 [19]. Additionally, in synovial sarcoma, the inclusion of an oncogenic fusion protein evicts a core subunit from the complex and leads to retargeting of the remainder complex [20]. However, currently no study has investigated the effect of removing one of the mutually exclusive subunits on the chromatin occupancy of its sibling subunit. Loss of a single member of the complex is the most frequent type of SWI/SNF alteration in cancer, therefore understanding how this affects the remaining complexes function in mechanistic detail is critical.

In this study we examine how loss of one subunit of a mutually exclusive pair affected the localization of its sibling subunit. We conducted an analysis on the two mutually exclusive ATPAse subunits, BRG1 and BRM (also known as SMARCA4 and SMARCA2, respectively). The functional interactions between BRG1 and BRM were not uniform. The predominant effect in HepG2 cells of loss of either BRG1 or BRM is the loss of the remaining subunit (~50-70% of sites). This occurs at genes that are both activated and repressed by BRG1 and BRM. In a subset of cases the sibling subunit is selectively retained or gained upon loss of the other subunit from chromatin (~30-50% of sites). A predominant loss of SWI/SNF occupancy upon loss of one subunit has been observed in other cell types [21]. Similar to our findings on ARID1A and either ARID1B or ARID2, BRG1 and BRM often antagonize one another in transcriptional control. In cases of antagonism, a combined knockdown of BRG1 and BRM rescued the expression changes globally. These results demonstrate that complex functional interactions exist between mutually exclusive SWI/SNF complexes and highlight the importance of characterizing the effects on the complexes that remain in SWI/SNF mutated tumors. We present a model to understand the heterogeneity of regulation, where interactions between distinct SWI/SNF complexes acts as a first layer of transcriptional control. Loss of one form of the complex unmasks a second layer of transcriptional control that can be either activating or repressive depending on the specific chromatin regulators recruited to a site.

## Results

### BRG1 and BRM depletion lead to distinct morphology changes in HepG2 cells

We investigated the effect of depleting one of a mutually exclusive pair of sibling SWI/SNF subunits on transcription and on SWI/SNF occupancy in HepG2 cells using a series of ChIP-seq and RNA-seq experiments. We selected HepG2 because they are a common and well studied liver cancer cell line and extensive ENCODE ChIP-seq and chromatin state data are available for these cells. Using shRNA targeting either BRG1 or BRM we depleted HepG2 cells of these proteins and performed RNA-seq or ChIP-seq in these models. Knockdown of BRG1 using two shRNAs in HepG2 cells causes cells to become elongated or spindly (Fig. 1A, 1C). Knockdown of BRM using two shRNAs led to a different morphology with cells appearing more flattened and lacking the tightly packed morphology of a non-silencing control (Fig. 1B, 1D). This suggested that BRG1 and BRM have distinct effects on HepG2 cells. We selected the stronger shRNA (BRG1 shRNA#2 and BRM shRNA#1) to perform further RNA-seq and ChIP-seq experiments.

**Figure 1:**
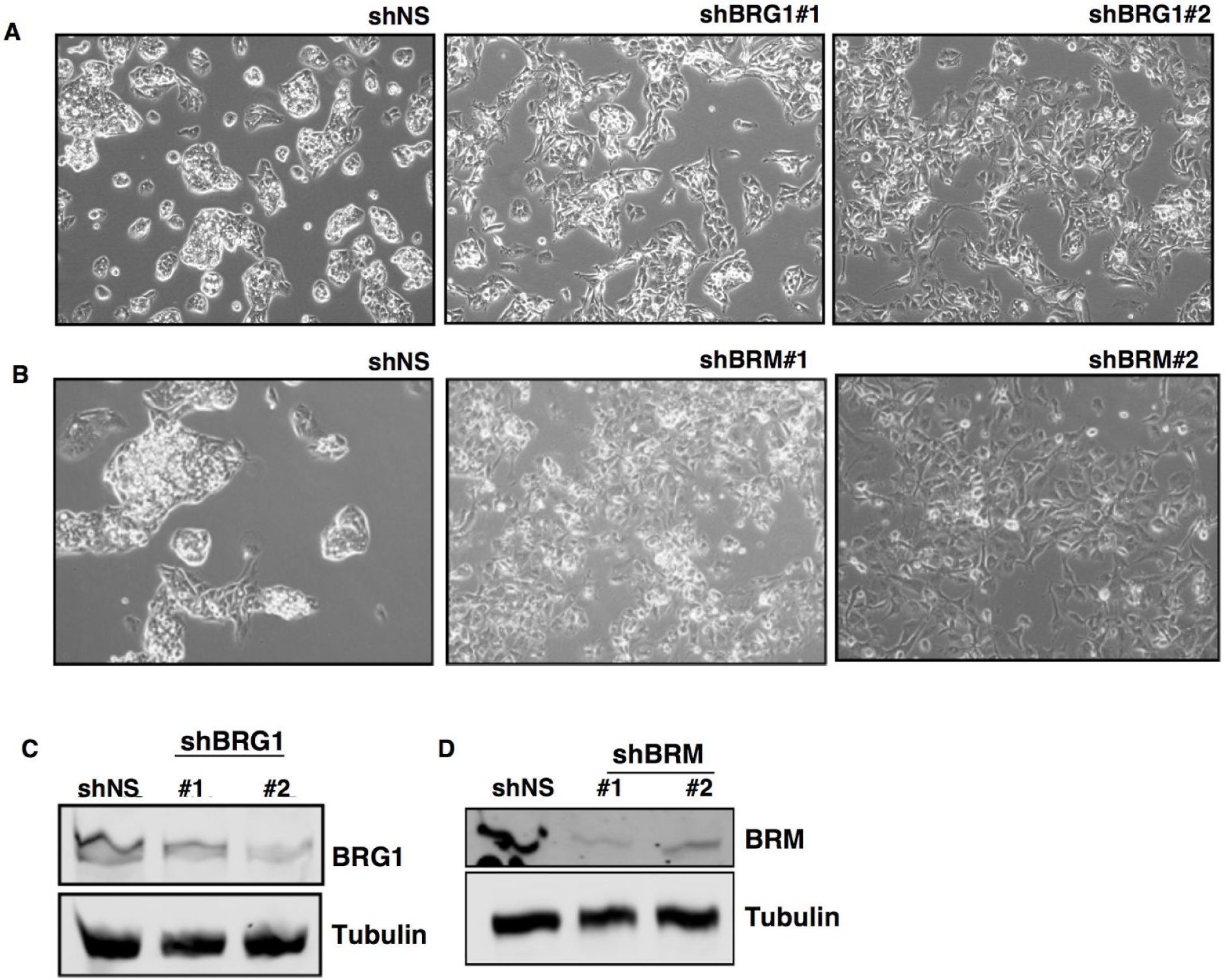
Depletion of BRG1 in HepG2. A. Morphology changes in HepG2 following treatment with two shRNAs targeting BRG1. B. Morphology changes in HepG2 following treatment with two shRNAs targeting BRM. C. Western blot of BRG1 following knockdown. D. Western blot following BRM knockdown.

### Sibling subunits control distinct and overlapping transcriptional programs

We first compared the effect of knockdown of BRG1 to BRM by RNA-seq. ShRNA targeting BRG1, BRM, or targeting both BRG1 and BRM were used to deplete HepG2 of the mutually exclusive subunits and subjected to RNA-seq. The shRNAs selectively targeted the appropriate subunit at the protein (Fig. 2A) and RNA level (Fig. 2B). At the RNA level most SWI/SNF subunits are unaffected by knockdown of either BRG1 or BRM or knockdown of both combined (Supplemental Fig. 1). Consistent with our observations in Figure 1, shBRG1 and shBRM induced distinct changes to cell morphology (Fig.2C). The double knockdown was a mixture of the two, although with considerably less severe morphology changes than shBRG1 alone.

**Figure 2:**
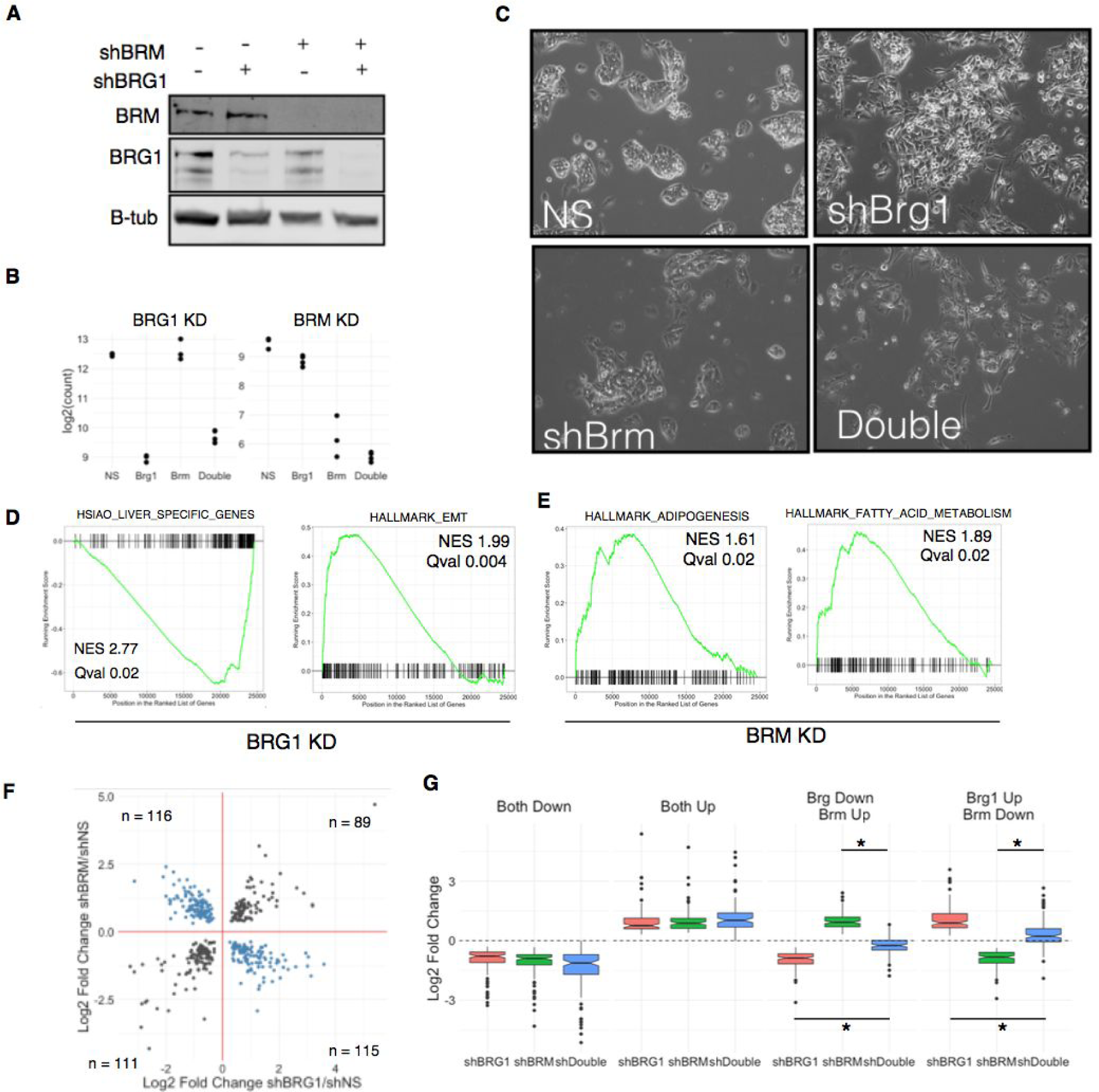
RNA-seq in BRG1, BRM, and combined knockdown. A. Western blot of HepG2 cells treated with shRNA targeting BRG1, BRM, or both combined. B. Log2 normalized read counts of RNA-seq data HepG2 treated with BRG1, BRM or both combined shRNA. C. Morphology changes of shRNA conditions. D. GSEA analysis of expression changes following shRNA targeting BRG1 and BRM. F. Fold changes of genes significantly differentially expressed and bound by BRG1 and BRM. G. Effect on genes in F following treatment with the individual and double knockdown shRNA.

We focused our analysis on genes that are direct targets of BRG1/BRM by analyzing genes associated with a ChIP-seq peak of BRG1 or BRM. To that end, we performed ChIP-seq in these models as well, and discuss the analysis of these data below (Fig. 3). For each peak we assigned the nearest gene as its direct target. We detected many direct target genes that changed following loss of BRG1 (3103), BRM (1188), or the double knockdown (3046) (Supplemental Table 1, 2, 3). We used GSEA analysis to identify categories of genes that are affected by BRG1 or BRM loss [22,23]. In the BRG1 knockdown we found a strong decrease in genes associated with liver identity and an upregulation of genes associated with an epithelial to mesenchymal transition (Fig.2D, Supplemental Table 4). Knockdown of BRM led to distinct changes in gene categories including an upregulation of genes involved in adipogenesis and fatty acid metabolism (Fig. 2E, Supplemental Table 5). In total, we found that 3860 genes were regulated by either BRG1 or BRM, and 431 genes were regulated by both BRG1 and BRM. Additionally, we observed 1023 directly regulated genes that were only affected in the double knockdown which may signify a redundant role for BRG1 and BRM in the regulation of these genes (19% of affected genes). In total, these data show that 5314 were dysregulated by loss of at least one SWI/SNF ATPase subunit of the total 35010 genes with measureable transcription in our assay (15%).

**Figure 3:**
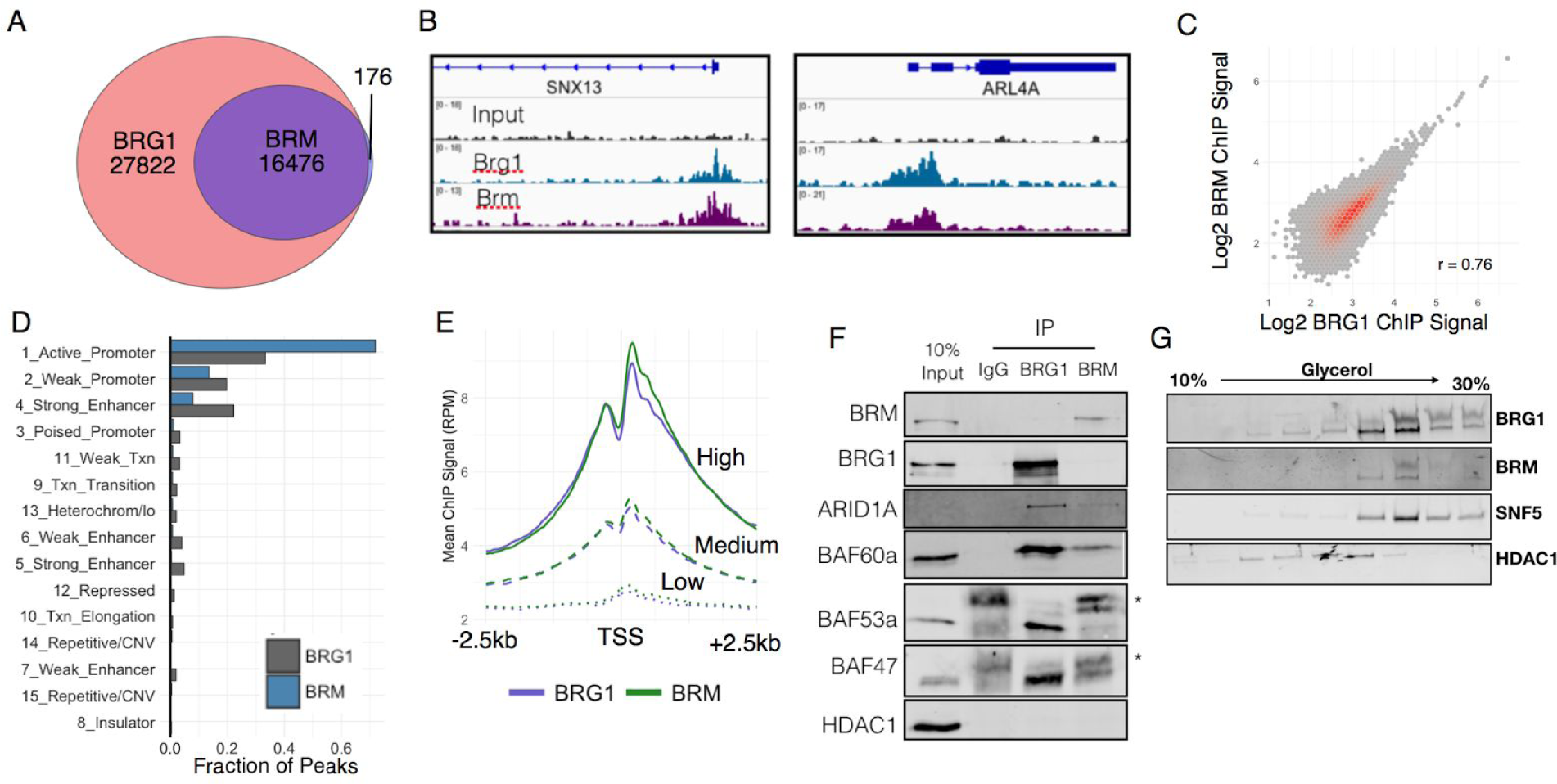
BRG1 and BRM occupy common genomic regions. A. Overlap between BRG1 and BRM ChIP-seq peaks. B. Example of BRG1 and BRM occupancy. C. Scatterplot of BRG1 and BRM signal at union set of BRG1 and BRM peaks. C. Venn diagram depicting overlap between BRG1 and BRM sites. D. ChromHMM features associated with BRG1 and BRM peaks. E. BRG1 and BRM ChIP-seq signal centered on TSS stratified by expression level of gene into low, medium, and high expression. F. Co-immunoprecipitation of BRG1 and BRM followed by western blot of SWI/SNF subunits. Asterisks denote IgG heavy chain. G. Glycerol ultracentrifugation of SWI/SNF subunits

Plotting the fold changes of shBRG1 against shBRM revealed that direct targets of BRG1 and BRM were both concordantly and discordantly regulated (Fig. 2F), similar to our previous observations for ARID1A and ARID1B [16]. We assigned each gene to a particular class based on the observed expression changes following BRG1 and BRM loss. This yielded four categories, Both Up (n = 89), Both Down (n = 111), BRG1 Up BRM Down (n = 115), BRG1 down BRM up (n = 116) (Fig. 2F, Supplemental Table 6). We then analyzed expression at each of these classes following knockdown by the single or double shRNA. At sites that were regulated concordantly (Both Up or Both Down), the double knockdown exaggerated the effect slightly, driving expression slightly higher or lower than the single knockdown (Fig. 2G). At genes that were regulated discordantly however there was a robust effect of the double knockdown. In both cases where BRG1 loss upregulated a gene or downregulated a gene and BRM had the opposite effect, the double knockdown strongly blunted these changes returning expression nearly to baseline (Fig. 2G). Therefore, at genes where BRG1 and BRM function regulate genes discordantly, we found they functionally antagonize one another. We previously observed a similar effect between ARID1A and either ARID1B or ARID2, but at only small number of genes tested individually. These data confirm our previous result and extend it to a new set of mutually exclusive subunits genome wide [16]. These data demonstrate that BRG1 and BRM can have both cooperative and antagonistic effects on gene expression.

### BRG1 and BRM concordant and discordant sites are associated with distinct transcriptional programs

We performed gene ontology analysis using categories from the Molecular Signature Perturbations Database (MSigDB) using the C2 (genetic and chemical perturbations) and the H (Hallmark) series of annotations ([22,23], Supplemental Table 6). Genes associated with concordant BRG1/BRM regulation that both increase following BRG1 or BRM loss were associated with TNF-alpha signaling through NFKB. Genes associated with concordant BRG1/BRM regulation that both decrease following BRG1/BRM loss were associated with genes upregulated in hepatoblastoma (CAIRO_HEPATOBLASTOMA_UP) or liver cancer with CTNNB1 mutations (CHIANG_LIVER_CANCER_SUBCLASS_UP). These categories are consistent with the cellular origin of HepG2 and might suggest expression of BRG1 and BRM were important for transformation of this tumor. In cases where BRG1/BRM regulated a gene discordantly, these genes were associated with several ontologies that discriminated between two subsets of cancer types (CHARAFE_BREAST_CANCER_LUMINAL_VS_BASAL_DN and CHARAFE_BREAST_CANCER_LUMINAL_VS_MESENCHYMAL_DN). Finally, genes specifically altered only when both BRG1 and BRM are depleted (redundant genes) were enriched for ontologies associated with general transcription, cell signaling, and apoptosis (REACTOME_GENERIC_TRANSCRIPTION_PATHWAY, REACTOME_CELL_CELL_COMMUNICATION, PID_PIK3CIPATHWAY, CONCANNON_APOPTOSIS_BY_EPOXOMICIN_UP - See Supplemental Table 7 for full results)

### BRG1 and BRM occupy similar genomic loci

We next wanted to determine how the localization of BRG1 and BRM relate to one another, therefore we performed a series of ChIP-seq experiments in shNS, shBRG1, and shBRM cells. In shNS cells there was a high degree of overlap between BRG1 and BRM. Nearly all BRM peaks overlapped a BRG1 peak (Fig. 3A). The high degree of similarity was evident at specific sites as well as genome wide (Fig. 3B, 3C). The lack of complete overlap between BRG1 and BRM is likely due to differences in antibody efficiency given the high correlation in signal. BRG1/BRM occupancy was mostly associated with active regulatory regions of the genome based on ChromHMM annotations including promoters and strong enhancers (Fig. 3D, Supplemental Table 8,9) [24]. The high degree of ChIP-seq overlap was not due to the antibodies precipitating both ATPase subunits together. Co-immunoprecipitation experiments show that BRG1 and BRM immunoprecipitated other SWI/SNF members (ARID1A, BAF60A, BAF53A, BAF47) while not immunoprecipitating each other (Fig. 3F). The IP of BRM was weaker than BRG1 due either to lower protein abundance or IP efficiency. Glycerol gradient ultracentrifugation analysis shows that both BRG1 and BRM are found in later fractions indicating both subunits assemble into similar sized SWI/SNF complexes (Fig. 3G). HDAC1 is not a SWI/SNF complex member and is not immunoprecipitated by either antibody and is found in a distinct set of fractions following ultracentrifugation on a glycerol gradient (Fig. 3F, 3G).

### Depletion of BRG or BRM leads to loss of occupancy of the remaining sibling complex

We next wished to determine how localization one sibling subunit is affected following loss of the other sibling. To address this we performed ChIP-seq on BRM and BRG1 following knockdown of either BRG1 or BRM (Supplemental Table 7, 8). Knockdown of BRG1 led to a decrease genome-wide of BRG1 signal to approximately half of shNS levels (Fig. 4A, 4B). Depletion of BRM led to a very robust decrease to near background levels of BRM occupancy (Fig. 4A, 4C). Consistent with the overall decrease in signal, we observed a reduction in the number BRG1 peaks following BRG1 knockdown from (44298 in shNS to 19032 in shBRG1, Fig. 4E, Supplemental Table 7). In shBRM treated HepG2 cells we identified only 90 peaks compared to 16652 in shNS cells, suggesting BRM is nearly completely depleted in these cells (Fig. 4F, Supplemental Table 8). The difference in efficiency could reflect difference in shRNA activity, differences in normal levels of these proteins, or the differences in the ability of cells to tolerate a complete loss of BRG compared to BRM.

**Figure 4:**
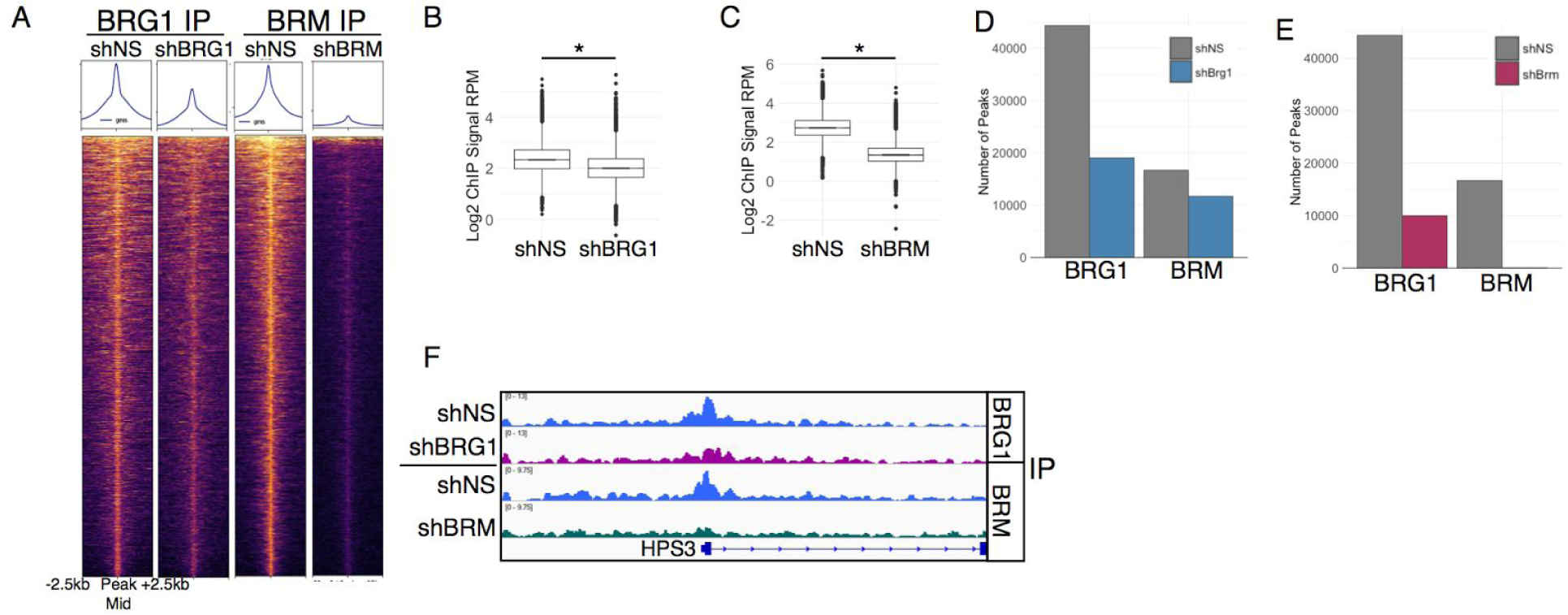
ChIP-seq on BRG1 and BRM in cells depleted of these subunits. A. Heatmap showing effect of BRG1 and BRM depletion on BRG1 and BRM occupancy at all BRG1 or BRM sites from non targeting treated cells. B,C Quantification of BRG1 (B) or BRM (C) signal from panel A. D, E. Number of peaks obtained in shBRG and shBRM treated cells for BRG1 and BRM ChIP experiments compared to shNS treated cells. F. Example locus showing moderate depletion of BRG1 and near complete loss of BRM following shRNA treatment.

We then compared the effect of shBRG1 and shBRM to the occupancy of the opposite ATPase. That is, how does loss of BRM affect BRG1 occupancy genome-wide (Fig. 5A-D). We focused our analysis on the the regions of the genome that lost occupancy for the first subunit because these are the most likely places to observe a robust change in the sibling subunit. At the 25913 regions that loss a BRG1 peak in shBRG1 treated cells we observed a moderate decrease in both BRG1 and BRM signal in shBRG1 (Fig. 5A, 5B). At these sites shBRM treatment led not only to the decrease in BRM occupancy but also led to a strong decrease in BRG1 occupancy. We performed the reverse analysis, focusing on the 16564 lost BRM peaks in shBRM treated cells. These sites also lost occupancy of the sibling subunit. For example, BRG1 ChIP-seq signal in shBRM treated cells decrease relative to shNS (Fig. 5C,D). Together, these data suggest that BRG1 and BRM reinforce one another's occupancy despite not physically interacting with one another. This was not due to off-target shRNA effects or cross reactivity between our antibodies (Fig. 2A, 2B, 3E). Despite mostly a decrease in signal, slightly less than half of sites overlapped a region that remained called as a peak or was a new peak. We classify these sites as retained.

**Figure 5:**
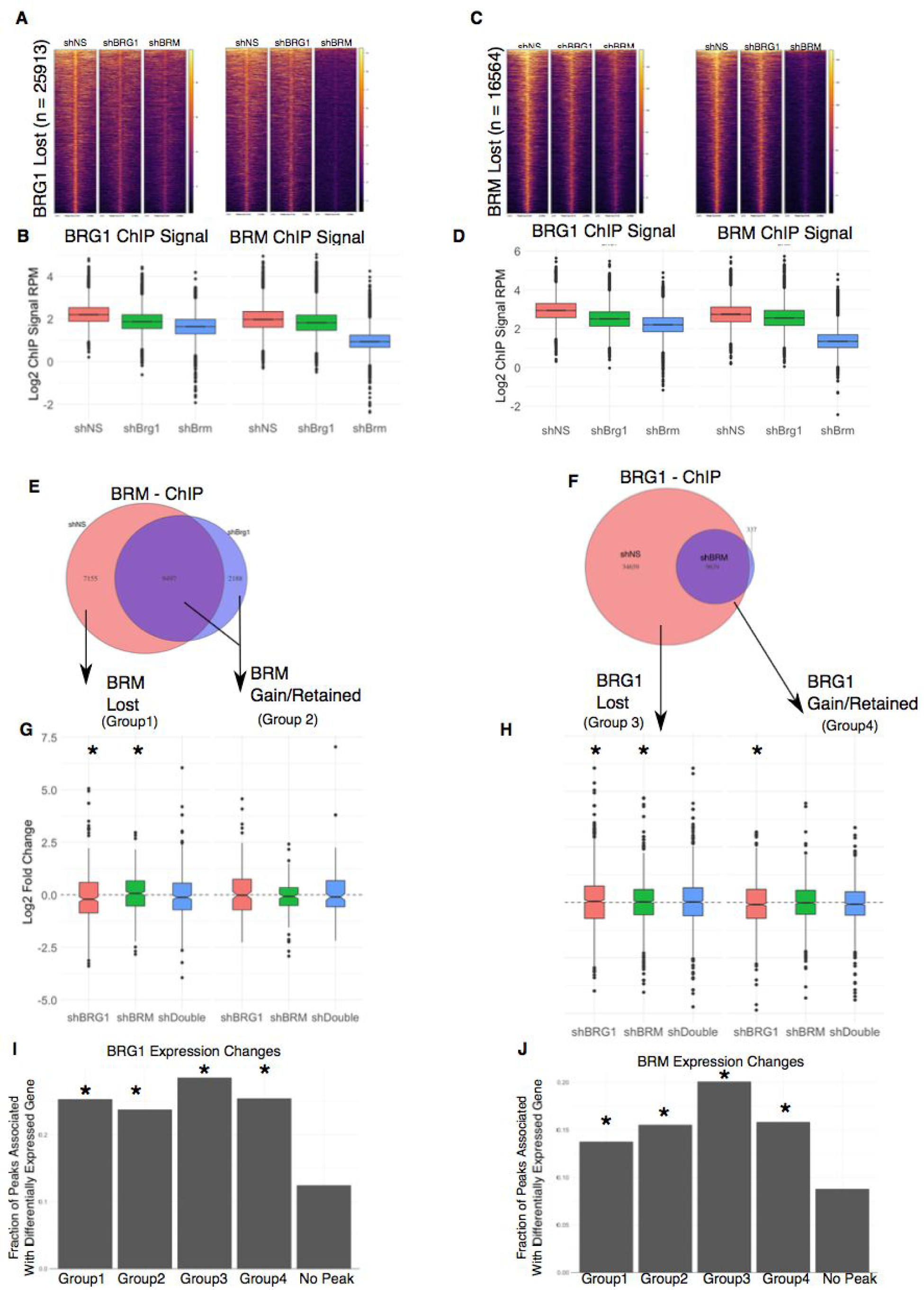
Occupancy and transcriptional changes associated with BRG1 or BRM loss. A-D. Effect of BRG1 or BRM depletion of BRG (A, B) or BRM (C, D). E, F. Classification of ChIP-seq peaks based on loss of BRG and BRM. G, H Quantification of gene expression changes in categories assigned in E,F. Asterisk denotes p-value < 0.05 by wilcox rank sum test against hypothesis of no change in expression. I, J. Fraction of peaks in each category associated with significantly altered gene expression. Asterisks denotes p-value < 0.05 by chi square test relative to genes that were not associated with a BRG/BRM altered peak.

### Loss of BRG1/BRM occupancy is associated with both activating and repressing activity

To determine how loss of or retention of the sibling subunit regulated transcription we grouped the genes associated with peaks that lose a subunit into four categories (Fig. 5E, 5F). Group1 genes are genes that lose BRG1 and lose a BRM peak. Group 2 genes are those that lose BRG1 and gain or retain a BRM peak. Group 3 genes are those that lose BRM and lose BRG1, and group 4 genes are those that lose BRM and gain or retain BRG1. We chose to classify gained/retained together because we could not be sure these reflected actual new peaks or quantitative differences in ChIP occupancy that increased above the threshold necessary for a a peak to be called. Using this classification scheme we analyzed the expression changes from our RNA-seq experiment and performed GSEA analysis. Many of the genes associated with each of the categories changed expression. There was a modest but significant difference at these categories in response to different shRNA conditions (Fig. 5G, 5H). For example, in group 1 genes, shBRG1 caused genes to decrease expression, while shBRM caused genes to increase expression (Wilcoxon test, p-value < 0.05 against hypothesis of no fold change). This effect was consistent with our observation that BRG1 and BRM could function antagonistically. Finally, we looked broadly at whether loss of BRG1/BRM peaks affected transcription of the associated genes without specifically asking about the direction or magnitude of change. Compared to genes that were not associated with a BRG1 or BRM peak in one of the above categories, genes in all four groups were more significantly more likely to affect gene expression (chi square test, p-value < 0.05).

We explored the functional annotations associated with these and again found that specific categories of genes were affected differentially by BRG1 and BRM (Supplemental Table 10). Among genes where BRG1 is lost, but BRM is gained or retained we found enrichment for genes involved in TNF-alpha signaling through NFKB (Fig. 6A). Notably, this enrichment was affected opposingly by BRG1 and BRM and is consistent with occupancy changes where BRG1 is lost and BRM remains. In cells treated with shBRG1, expression of this gene set increased showing that BRG1 is a negative regulator of this pathways transcription while BRM is a positive regulator. A second example was found in group four. Here we found that BRG1 negatively regulated fatty acid metabolism, while BRM was the positive regulator. These results suggest that BRG1 and BRM act as site specific regulators of transcription, and the direction of that regulation cannot be explained simply by their occupancy.

**Figure 6:**
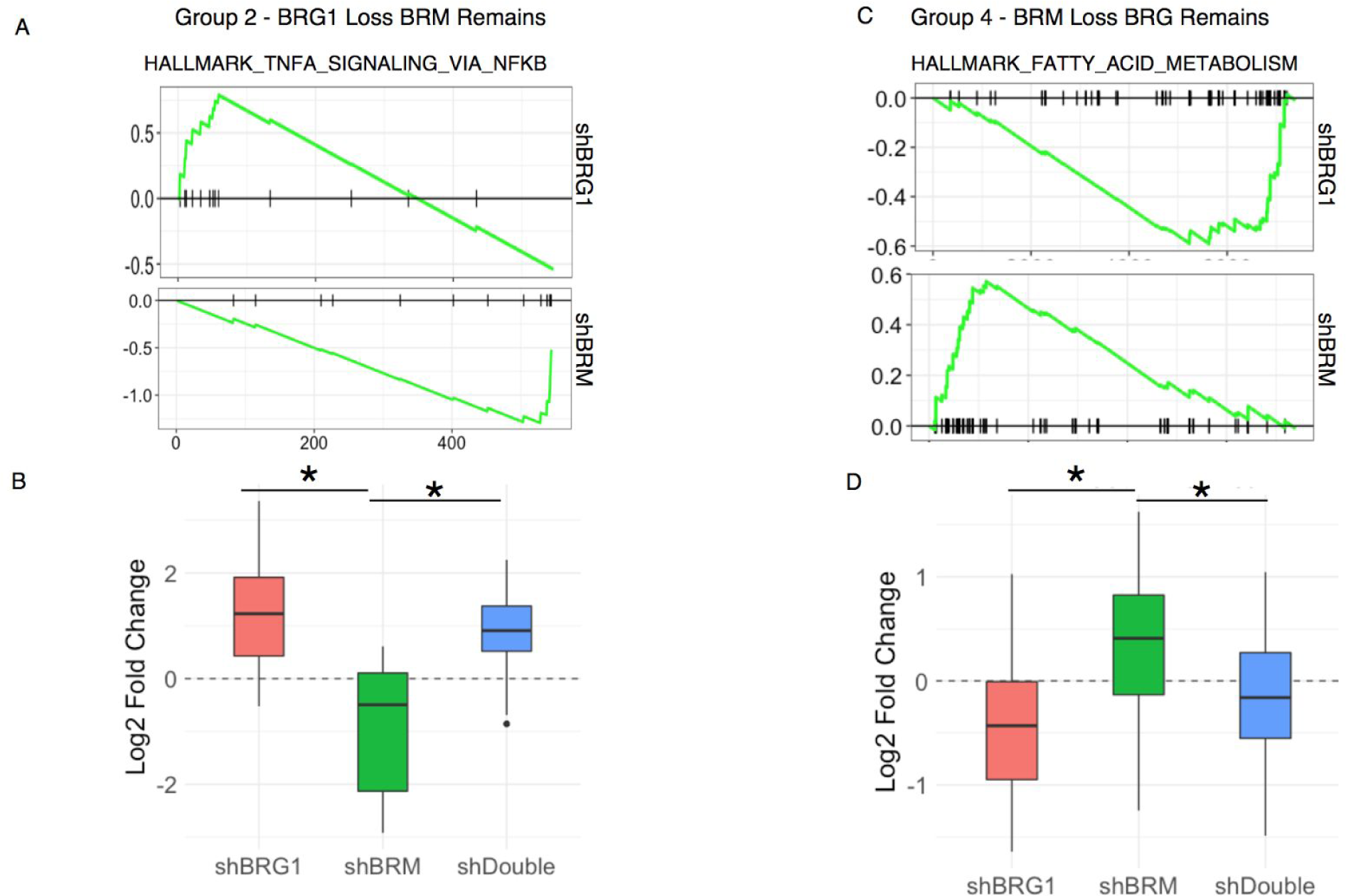
Different regulatory classes are composed of distinct gene sets. A. GSEA analysis of group 2 genes (BRG1 loss, BRM remains) showing different effects of shBRG1 compared to shBRM on expression in this gene signature. B. Quantification of gene expression changes in the Hallmark TNF-alpha Signaling through NFKB group. C. GSEA analysis of Group4 (BRM loss, BRG remains) showing different effects of shBRG1 compared to shBRM on a gene signature associated with fatty acid metabolism. D. Quantification of C.

### Altered SWI/SNF transcriptional controls is associated with distinct cofactors

Our results demonstrate that BRG1 and BRM have numerous functional interactions in HepG2 cells. The effect on transcription of these interactions does not depend simply on the occupancy of BRG1 and BRM, but through some site-specific regulatory control. One possibility is that BRG and BRM may directly or indirectly interact with other chromatin and transcriptional regulators to control the output at a specific site. To address this hypothesis we made use of the extensive data available from the ENCODE project for HepG2. Using these data we compared the level of ChIP-seq signal for each factor available in ENCODE at regions that lose both sibling subunits to those that lose one sibling while retaining or gaining the other sibling. We did not find any factors for which there was complete absence in one group and presence in another. However, we found several examples of factors with quantitative differences in levels (Fig. 7A-7C). We found a strong enrichment for EZH2 at regions that lose both BRG1 and BRM (Fig. 7A). Conversely, we found that HNF4A and FOXA1 were more strongly enriched at regions that lost BRM but gain or retain BRG1(Fig. 7B, 7C). These data suggest that functional interactions between different SWI/SNF complexes and other chromatin regulating complexes may play a role in the ultimate outcome of SWI/SNF disruption.

**Figure 7:**
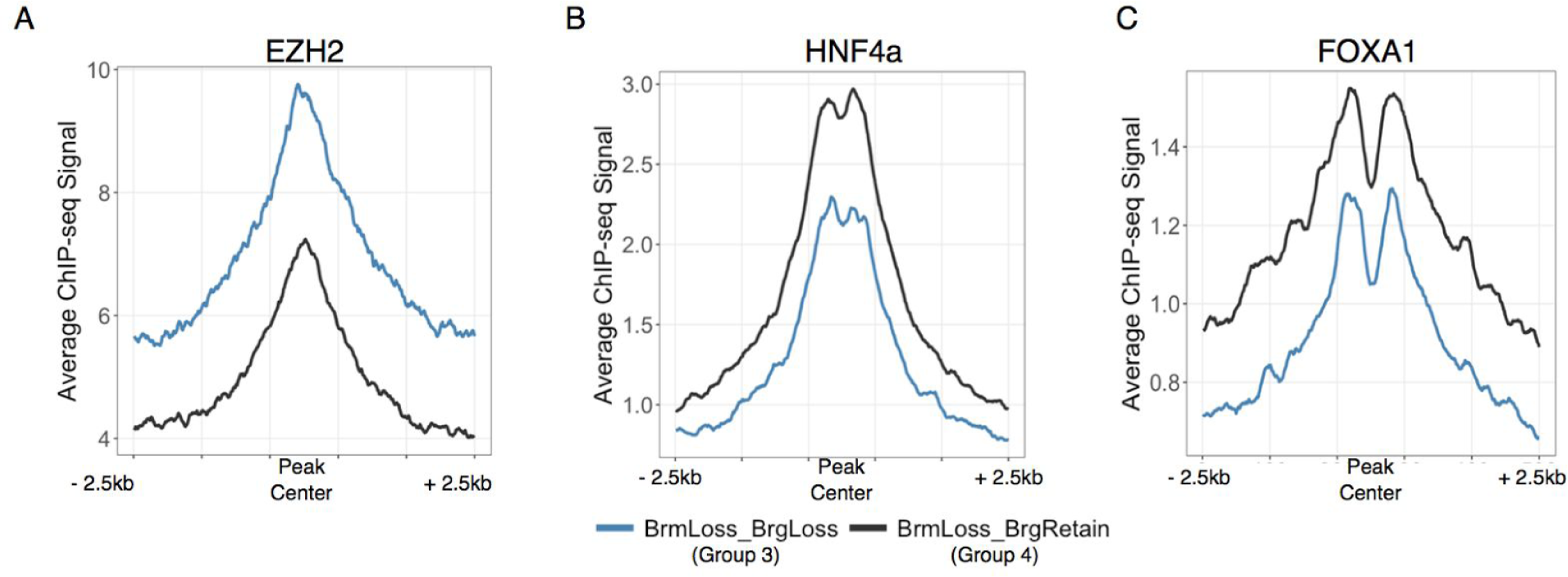
Distinct co-factors are associated with lost compared to retained/gained sites. A-C ChIP-seq signal for EZH2 (A) HNF4a (B), and FOXA1 (C) aligned to the midpoint of BRM peaks that following BRM depletion either lose (Group 3) or retain/gain (Group 4) BRG1.

## Discussion

In this study we aimed to understand how occupancy by two mutually exclusive SWI/SNF complexes, distinguished here by the ATPase subunit, changed when one form of the complex was depleted. Our goal was to tie these changes in occupancy to changes in transcription and identify factors that contribute to the choice of mechanism. We found that transcriptional control and occupancy are complex and include a myriad of mechanisms that are controlled in site-specific ways (Fig. 8). Careful dissection of the factors that associate differentially with BRG1 and BRM are required for elucidating how a specific site is targeted by different mechanisms. Our study highlights that SWI/SNF complexes are both biochemically and functionally heterogenous and genome-wide assessments of their role in transcriptional regulation mediated by multiple subunits are needed to gain an accurate picture of their effect in a biological system of interest.

**Figure 8:**
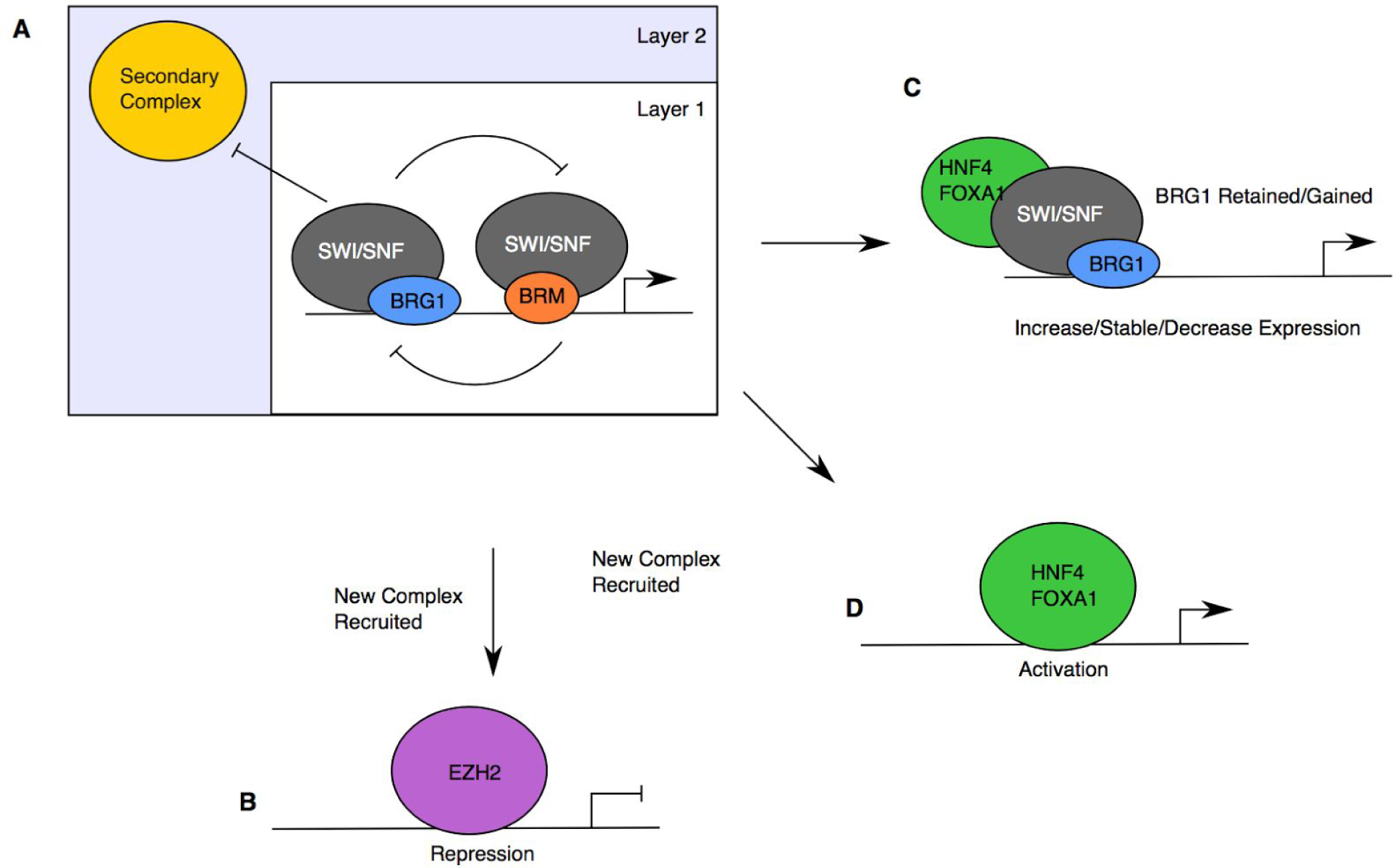
Hierarchical model of functional interactions between BRG1 and BRM. A. Interactions in layer 1 are primarily between SWI/SNF complexes, excluding secondary complexes form the locus. Upon loss of one of the ATPase subunits, the remaining complex association with the locus, and transcriptional output is controlled by the identity and action of the secondary complex. Interactions with the remaining SWI/SNF complex supplant SWI/SNF occupancy (B,D) or retain it C). If the the remaining complex is evicted, the transcriptional output is controlled by the whether the new complex is repressive (B) or activating (C,D).

### Functional interactions control SWI/SNF occupancy and transcriptional output

At the occupancy level, we find primarily that BRG1 and BRM bind commonly throughout the genome. The level of BRG1 and BRM correlates well with overall transcription. At approximately half of sites depletion of one subunit leads to loss of the second subunit (70% for BRM, 40% for BRG1, suggesting a cooperative force in binding. However, at a subset of sites, the sibling subunit is gained or may increase in occupancy.

The mechanisms of transcriptional regulation mediated by mutually exclusive subunits of SWI/SNF has previously been grouped into two categories. In one, studies show that BRG1 and BRM are redundant, and the loss of one subunit unmasks a requirement or a new function for the other subunit. This principle underlies the observed synthetic lethality seen between BRG1 and BRM in multiple cancers [25,26]. Similar mechanisms are believed to cause the synthetic lethality observed between ARID1A and ARID1B [27]. Our study provides evidence of this from the 1023 genes specifically affected following knockdown of both BRG1 and BRM (~20% of genes affected).

A second category of regulatory mechanism is antagonism. In this case the two mutually exclusive subunits have opposing effects on transcription. This has been shown in a developmental context and at specific genes [4,28]. We previously observed strong antagonism between ARID1A and ARID1B, as well as between ARID1A and ARID2 at a subset of genes in HepG2 cells. Through double knockdown experiments at a small number of genes we showed that the changes in expression observed following knockdown of ARID1A depended on ARID1B and ARID2 [16]. Our expression and occupancy data demonstrate that this mechanism is also found for BRG1/BRM and suggests it is generally applicable to mutually exclusive subunits. We find a group of genes where BRG1 plays an activating role in transcription while BRM plays a repressive role. We observe another set of genes where the reverse is true, suggesting even in a single cell type there is not a consistent functional output to co-occupancy and co-regulation by BRG1 and BRM.

We observe a third class where BRG1 and BRM are cooperative. At these genes loss of BRG1 was equivalent to the loss of BRM and the reverse was true. The linkage between cooperative occupancy and cooperative expression regulation was not one-to-one. The predominant effect on occupancy appeared cooperative, even in instances where this did not lead to concordant gene expression changes.

### A model linking transcription output and SWI/SNF occupancy

The complexity of these interactions make defining a single model difficult for SWI/SNF action. We propose one possible model for how this occurs that involves hierarchical interactions between chromatin regulators (Fig. 8A). The first layer of the model is dependent on interactions between BRG1 containing SWI/SNF complexes and BRM containing SWI/SNF complexes. In this model, BRG1 and BRM associate with their own cofactors to regulate transcription. Such interactions can be cooperative or competitive, in both transcription and occupancy, depending on the specific accessory complexes involved. The baseline expression of a locus under this condition is ‘On’, but the precise level is controlled by the interaction between the two SWI/SNF complexes. If both SWI/SNF complexes are positive regulators, expression will be high. If the SWI/SNF complexes antagonize one another, the level of expression will be intermediate and tuned by the functional interaction between SWI/SNF complexes.

The second layer of regulation of the locus may involve a second chromatin regulating complex. At baseline, the second complex is not sufficient to evict both SWI/SNF complexes, and SWI/SNF regulation dominates at the locus. Upon loss of one of the ATPase subunits, the remaining subunit is also lost because a repressive complex can now dominate and evict the second SWI/SNF complex. In a population of cells, one would expect to identify some enrichment for this second complex, indicative of a fraction of the cell population where the secondary complex is functional.

This model is consistent with the various transcriptional outputs that are observed following disruption of SWI/SNF occupancy. If the second layer of regulation is controlled by a repressive complex, such as PRC2, then it may evict SWI/SNF entirely, leading to both decreased occupancy and decreased transcription (Fig. 8B). If the second layer of regulation is by a positive regulator, such as HNF4A or FOXA1, the sibling subunit may be evicted if the secondary complex has its own strong chromatin regulator associated (Fig. 8C). This would lead to complete loss of SWI/SNF, but may increase transcription. Alternatively, the secondary complex may be capable of interacting with the remaining sibling subunit. This could lead to retention and a transcriptional output that is dependent on the initial state (Fig. 8D).

The underlying mechanism that controls how a particular site is regulated remains unclear. Our analysis of ENCODE factors identified chromatin regulators and transcription factors that were specific to sites that either lose both subunits or retain one of the sibling subunits. This suggests that at least functional interactions between SWI/SNF and other chromatin regulators are involved [29–32]. A second possibility is that post-translational modifications of SWI/SNF or associated regulators control assembly and transcription at these sites. Few studies have looked at these modifications on SWI/SNF, however in breast cancer, methylation of BAF155 by CARM1 is important for SWI/SNF activity in metastatic progression [33]. A final possibility is that the composition of SWI/SNF differs between the different classes of sites. Future studies aimed at biochemically purifying subsets of SWI/SNF based on post-translational modifications or genomic loci are needed to determine which mechanisms above are important.

## Methods

### Cell culture

HepG2 cells were purchased from ATCC. They were grown in DMEM (Gibco, Life Technologies) containing 10% fetal calf serum supplemented with 100units/mL penicillin/streptomycin (Life Technologies). Cells tested negative for mycoplasma contamination and were not passaged more than 6 times from initial cells from ATCC.

### Chip-seq

Cells were transduced with shRNA targeting BRG1 or a non-targeting control in the presence of 8ug/mL polybrene. 2ug/mL was added 24 hours post transfection, and cells were grown for a total of 7 days following transduction before harvest. ChIP was performed as previously described with modifications [16]. Cells were fixed for 30 minutes at 4C in 0.3% methanol free formaldehyde, quenched for 5 minutes with 125mM glycine, washed 3 times and snap frozen in liquid nitrogen and stored at −80C. Frozen pellets were thawed for 30 minutes on ice, resuspended each pellet in 1mL swelling buffer (25mM HEPES + 1.5mM MgCl2 + 10mM KCl + 0.1% Igepal containing 1mM PMSF and 1X protease inhibitor cocktail, Roche) and incubated 10 minutes at 4C. Cells were dounced 20 strokes with a “B” pestle and then nuclei were pelleted at 2000 rpm for 7 minutes at 4C. The nuclei were washed with 10mL MNase Digestion Buffer (15mM Hepes pH 7.9, 60mM KCl, 15mM NaCl, 0.32M Sucrose) and pelleted at 2000 rpm for 7 minutes at 4C. The pellet was then resuspended in 1mL MNase Digestion Buffer per 4e7 cells + 3.3uL 1M CaCl2 per mL + PMSF (1mM) and Protease Inhibitor Cocktail (1X, Roche), then incubated for 5 minutes at 37C to warm. We added MNase (NEB M0247S 2000U/ul) at 0.5uL/1e7 cells and incubated for 15 minutes at 37C with agitation. Following digestion, the MNase was chelated using 1/50 volume 0.5M EGTA on ice for 5 minutes. We added 1 volume of 2X IP Buffer (20mM TrisCl pH 8, 200mM NaCl, 1mM EDTA, 0.5mM EGTA), then passed the sample for a 21G needle 5 times and added Triton X-100 to 1% final concentration. The sample was cleared at 13,000 RPM for 15 minutes at 4 degrees and chromatin (30ug BRG1 (abcam ab110641) /BRM (Cell signaling 11966)) was used incubated with antibody overnight at 4 degrees. Antibody/chromatin complexes were captured with protein A magnetic beads (Bio-Rad) for 2 hours at 4 degrees, washed 5 times with Agilent RIPA (50mM Hepes pH 7.9/500mM LiCl/1mM EDTA/1% IGEPAL-ca-630/0.7% Na-Deoxycholate) and once with 10mM Tris/1mM EDTA/50mM NaCl. DNA was eluted at 65C with agitation using 100uL 1% SDS + 100mM NaHCO3 made freshly. Crosslinks were reversed overnight by adding 5uL of 5M NaCl and incubating at 65C. DNA was treated with 3uL RNaseA for 30 minutes at 37C and then Proteinase K for 1 hours at 56C and purified with Zymo Clean and Concentrator ChIP Kit and quantified using qubit before library preparation (Kapa Hyperprep).

## ChIP-Seq Data Analysis

### Mapping

Reads were aligned to hg19 using bowtie2 [34] using the ‐‐sensitive parameters, and duplicates were removed using samtools [35]. For visualization bigwig tracks were generated using Deeptools [36] (version 2.2.2) bamCoverage tool with a binsize of 10bp and extending fragments to the approximate nucleosome size of 150bp. Tracks can be visualized using IGV [37] and bigwig files are available in GEO Accession number GSE102559.

### Peak Calling

Peaks were called using Macs2 (version 2.1.0 [38]) using the narrowpeak mode using the following parameters. Qval = 0.01 ‐‐keep-dup-all ‐‐shift 37 ‐‐nomodel ‐‐extsize 147. Additionally, we filtered the peaks against the ENCODE blacklist regions and further recursively merged any peaks within 500bp of the nearest peak. Peak calls for each of the six data sets can be found in Supplemental Table 3.

### Gain vs Lost peaks

Using peak calls described above, we overlapped the peaks from shNS and shBRG1 or shBRM. We selected peaks that were lost called in shNS, but not called in on of the ATPase knockdown experiments to categorise peaks for further analysis. These were further grouped based on overlap of peaks in the opposite siblings experiment. For example, if a BRG1 peaks was lost between shNS and shBRG1, these BRM peak calls in these regions were overlapped in shNS and shBRG1 to identify BRM lost and BRM retained/gained peaks.

## RNA-seq

Cells were transduced with shRNA targeting BRG1, BRM, Both combined, or a non-targeting control in the presence of 8ug/mL polybrene. 24 hours after transduction, 2ug/mL puromycin was added and cells were maintained under selection until harvest (6 days). Cells were harvested in trizol and RNA was extracted using the Directzol RNA kit (Zymo). Libraries were prepared using the Kapa mRNA library kit per manufacturer's instructions and sequenced on a Hiseq 4000 (50bp).

## RNA-seq Data Analysis

Libraries were sequenced on Hiseq 4000 (50bp reads) Gene expression levels were quantitated using kallisto[39]. These data were converted to counts and summarized per gene using tximport [40] and differential expression was carried out using DESeq2[41] using an FDR of 0.05 and no explicit fold change cutoff. Raw count matrices used as input to DESeq2 and output of DESeq2 can be found in Supplemental Tables 1 and 2.

## Preparation of Nuclear Lysates

Cells were washed with PBS, then centrifuged at 1300 rpm for 10 min at 4C. Cells were washed with 20 packed cell volumes with hypotonic cell lysis buffer (10mM HEPES-KOH pH 7.9, 1.5mM MgCl2, 10mM KCl, 0.5mM DTT plus protease inhibitors) and placed on ice for 10 min to swell. Cells were then centrifuged at 1300 rpm for 10 min at 4C. Cells were dounced with B pestle in 2 packed cell volumes of hypotonic cell lysis buffer. Nuclei were pelleted at 1300 rpm for 10min at 4C, washed with 10 packed cell volumes with hypotonic cell lysis buffer and centrifuged at 5000 rpm for 10 min. Extractions were performed twice with 0.6 volume nuclear lysis buffer (20mM HEPES-KOH pH 7.9, 25% glycerol, 420 mM KCl, 1.5mM MgCl2, 0.2mM EDTA, 0.5mM and protease inhibitors). Lysates were clarified at 14,000 rpm for 10 min at 4C between extractions. Lysates were diluted with storage buffer (20mM HEPES-KOH pH 7.9, 20% glycerol, 0.2mM EDTA, 0.2mM DTT) to bring final KCl concentration to 150mM and stored at −80.

## Immunoaffinity

Prior to beginning the IP, we washed Dynal Protein A beads 3 times with PBS + 0.5% BSA at 4C. We resuspended beads in 400uL of 1X + 0.5% BSA, then added 4-10ug of antibody and incubated overnight at 4C. The following day, we thawed the nuclear lysates on ice. Lysates were added to antibody-conjugated beads and incubated overnight. Beads were washed 4 times with BC-150 (20mM HEPES-KOH pH 7.9, 0.15M KCL, 10% glycerol, 0.2mM EDTA ph 8.0, 0.1% Tween-20, 0.5mM DTT, and protease inhibitors), 2 x BC-100 (20mM HEPES-KOH pH 7.9, 0.1M KCL, 10% glycerol, 0.2mM EDTA ph 8.0, 0.1% Tween-20, 0.5mM DTT, and protease inhibitors), 1 x BC-60 (20mM HEPES-KOH, pH 7.9, 60mM KCl, 10% glycerol, 0.5mM DTT, and protease inhibitors). Proteins were eluted from beads using 2X Laemmli buffer with 100mM DTT for 10 minutes at 95C.

## Glycerol Gradient Centrifugation

Glycerol gradient centrifugation was performed as previously described [42]. Briefly, 500ug of nuclear lysate was loaded onto a 10-30% glycerol gradient and centrifuged at 40,000 RPM in SW-41 rotor for 16 hours at 4 degrees. 0.5mL fractions were collected and used for western blot analysis.

## Data Availability

Sequencing from this study is available under the SuperSeries GSE102561. ChIP-seq data is available in GSE102559. Expression data is available in GSE102560. Output for differential expression for the three conditions are found in Supplemental Table 1-3. GSEA analysis for BRG1 and BRM are in Supplemental Table 4-5. Expression values underlying Figure 2F, 2G are found in Supplemental Table 6. GSEA analysis for categorized RNA-seq changes (Figure 2F, 2G) are in Supplemental Table 7. Annotated peak calls from ChIP-seq experiments are found in Supplemental Table 8 (BRG1) and 9 (BRM). Supplemental Table 10 contains GSEA analysis of genes associated with Group 1 - Group 4 ChIP-seq peaks (Fig. 5). ENCODE data sets used. HNF4A - ENCFF267OJD. FOXA1 - ENCFF871KVO, EZH2 - ENCFF545PYA and ENCFF978OOP. Where two files are included, BAM files were merged into a single file.

## Declarations

### Author Contributions

Conceptualization, JRR and TM; Methodology, JRR; Investigation, JRR, JSR, CC; Data Curation, JRR; Writing – Original Draft, JR and TM; Writing – Review & Editing, JRR and TM; Funding Acquisition, JRR, and TM; Supervision, JRR and TM.

### Competing Interests

The authors declare no competing interests

### Consent for Publication

All authors read and approved this manuscript.

---

**Supplemental Figure 1:**
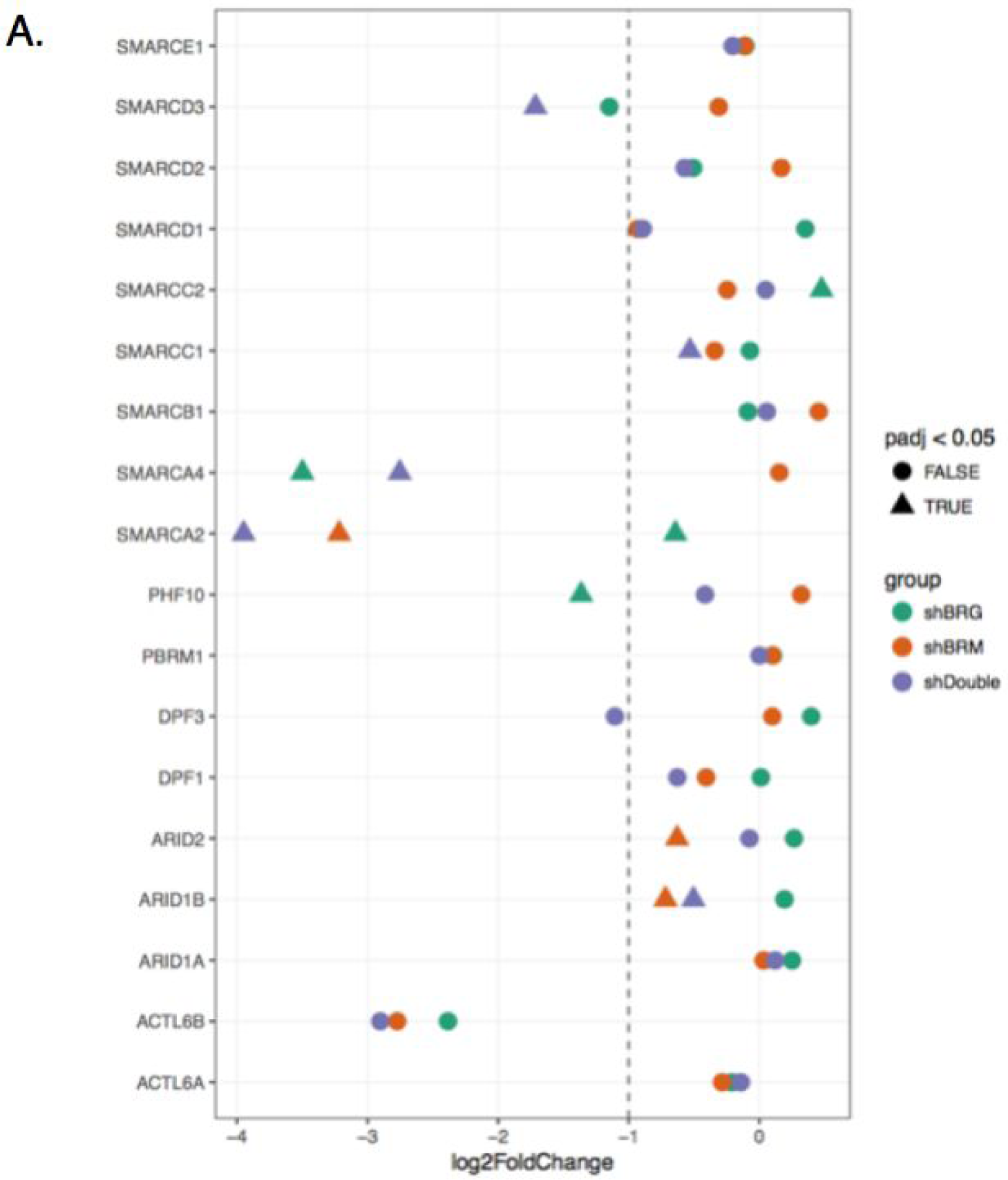
Expression changes of SWI/SNF subunits in RNA-seq experiments. Expression changes for each swi/snf subunit in the three RNA-seq knockdown conditions. SMARCA2 (BRM) and SMARCA4 (BRG1) are specifically downregulated in the appropriate shRNA treatment. Significant expression changes are denoted by triangles, and the color of the point denotes the shRNA treatment used.

## References

1. Clapier CR, Cairns BR. The biology of chromatin remodeling complexes. Annu. Rev. Biochem. 2009;78:273–304.

2. Hargreaves DC, Crabtree GR. ATP-dependent chromatin remodeling: genetics, genomics and mechanisms. Cell Res. Shanghai Institutes for Biological Sciences, Chinese Academy of Sciences; 2011;21:396–420.

3. Hodges C, Kirkland JG, Crabtree GR. The Many Roles of BAF (mSWI/SNF) and PBAF Complexes in Cancer. Cold Spring Harb. Perspect. Med. [Internet]. 2016;6. Available from: http://dx.doi.org/10.1101/cshperspect.a026930

4. Lessard J, Wu JI, Ranish JA, Wan M, Winslow MM, Staahl BT, et al. An essential switch in subunit composition of a chromatin remodeling complex during neural development. Neuron. 2007;55:201–15.

5. Flowers S, Nagl NGJr, Beck GRJr, Moran E. Antagonistic roles for BRM and BRG1 SWI/SNF complexes in differentiation. J. Biol. Chem. 2009;284:10067–75.

6. Shain AH, Pollack JR. The spectrum of SWI/SNF mutations, ubiquitous in human cancers. Kashanchi F, editor. PLoS One. Public Library of Science; 2013;8:e55119.

7. Kadoch C, Hargreaves DC, Hodges C, Elias L, Ho L, Ranish J, et al. Proteomic and bioinformatic analysis of mammalian SWI/SNF complexes identifies extensive roles in human malignancy. Nat. Genet. 2013;45:592–601.

8. Fujimoto A, Totoki Y, Abe T, Boroevich KA, Hosoda F, Nguyen HH, et al. Whole-genome sequencing of liver cancers identifies etiological influences on mutation patterns and recurrent mutations in chromatin regulators. Nat. Genet. Nature Publishing Group, a division of Macmillan Publishers Limited. All Rights Reserved.; 2012;44:760–4.

9. Guichard C, Amaddeo G, Imbeaud S, Ladeiro Y, Pelletier L, Maad IB, et al. Integrated analysis of somatic mutations and focal copy-number changes identifies key genes and pathways in hepatocellular carcinoma. Nat. Genet. Nature Publishing Group, a division of Macmillan Publishers Limited. All Rights Reserved.; 2012;44:694–8.

10. Li M, Zhao H, Zhang X, Wood LD, Anders RA, Choti MA, et al. Inactivating mutations of the chromatin remodeling gene ARID2 in hepatocellular carcinoma. Nat. Genet. Nature Publishing Group, a division of Macmillan Publishers Limited. All Rights Reserved.; 2011;43:828–9.

11. Le Loarer F, Watson S, Pierron G, de Montpreville VT, Ballet S, Firmin N, et al. SMARCA4 inactivation defines a group of undifferentiated thoracic malignancies transcriptionally related to BAF-deficient sarcomas. Nat. Genet. Nature Publishing Group, a division of Macmillan Publishers Limited. All Rights Reserved.; 2015;47:1200–5.

12. Versteege I, Sévenet N, Lange J, Rousseau-Merck MF, Ambros P, Handgretinger R, et al. Truncating mutations of hSNF5/INI1 in aggressive paediatric cancer. Nature. 1998;394:203–6.

13. Chandler RL, Damrauer JS, Raab JR, Schisler JC, Wilkerson MD, Didion JP, et al. Coexistent ARID1A-PIK3CA mutations promote ovarian clear-cell tumorigenesis through pro-tumorigenic inflammatory cytokine signalling. Nat. Commun. Nature Publishing Group; 2015;6:6118.

14. Jones S, Wang T-L, Shih I-M, Mao T-L, Nakayama K, Roden R, et al. Frequent Mutations of Chromatin Remodeling Gene ARID1A in Ovarian Clear Cell Carcinoma. Science. American Association for the Advancement of Science; 2010;330:228–31.

15. Wiegand KC, Shah SP, Al-Agha OM, Zhao Y, Tse K, Zeng T, et al. ARID1A mutations in endometriosis-associated ovarian carcinomas. N. Engl. J. Med. 2010;363:1532–43.

16. Raab JR, Resnick S, Magnuson T. Genome-Wide Transcriptional Regulation Mediated by Biochemically Distinct SWI/SNF Complexes. PLoS Genet. 2015;11:e1005748.

17. Kim KH, Kim W, Howard TP, Vazquez F, Tsherniak A, Wu JN, et al. SWI/SNF-mutant cancers depend on catalytic and non-catalytic activity of EZH2. Nat. Med. Nature Publishing Group, a division of Macmillan Publishers Limited. All Rights Reserved.; 2015;21:1491–6.

18. Wilson BG, Wang X, Shen X, McKenna ES, Lemieux ME, Cho Y-J, et al. Epigenetic antagonism between polycomb and SWI/SNF complexes during oncogenic transformation. Cancer Cell. 2010;18:316–28.

19. Stanton BZ, Hodges C, Calarco JP, Braun SMG, Ku WL, Kadoch C, et al. Smarca4 ATPase mutations disrupt direct eviction of PRC1 from chromatin. Nat. Genet. [Internet]. Nature Research; 2016 [cited 2016 Dec 12]; Available from: http://dx.doi.org/10.1038/ng.3735

20. Kadoch C, Crabtree GR. Reversible disruption of mSWI/SNF (BAF) complexes by the SS18-SSX oncogenic fusion in synovial sarcoma. Cell. 2013;153:71–85.

21. Mathur R, Alver BH, San Roman AK, Wilson BG, Wang X, Agoston AT, et al. ARID1A loss impairs enhancer-mediated gene regulation and drives colon cancer in mice. Nat. Genet. [Internet]. Nature Research; 2016 [cited 2016 Dec 12]; Available from: http://dx.doi.org/10.1038/ng.3744

22. Liberzon A, Subramanian A, Pinchback R, Thorvaldsdóttir H, Tamayo P, Mesirov JP. Molecular signatures database (MSigDB) 3.0. Bioinformatics. 2011;27:1739–40.

23. Liberzon A, Birger C, Thorvaldsdóttir H, Ghandi M, Mesirov JP, Tamayo P. The Molecular Signatures Database (MSigDB) hallmark gene set collection. Cell Syst. 2015;1:417–25.

24. Ernst J, Kheradpour P, Mikkelsen TS, Shoresh N, Ward LD, Epstein CB, et al. Mapping and analysis of chromatin state dynamics in nine human cell types. Nature. Nature Publishing Group, a division of Macmillan Publishers Limited. All Rights Reserved.; 2011;473:43–9.

25. McDonald ER 3rd, de Weck A, Schlabach MR, Billy E, Mavrakis KJ, Hoffman GR, et al. Project DRIVE: A Compendium of Cancer Dependencies and Synthetic Lethal Relationships Uncovered by Large-Scale, Deep RNAi Screening. Cell. 2017;170:577–92.e10.

26. Wilson BG, Helming KC, Wang X, Kim Y, Vazquez F, Jagani Z, et al. Residual complexes containing SMARCA2 (BRM) underlie the oncogenic drive of SMARCA4 (BRG1) mutation. Mol. Cell. Biol. 2014;34:1136–44.

27. Helming KC, Wang X, Wilson BG, Vazquez F, Haswell JR, Manchester HE, et al. ARID1B is a specific vulnerability in ARID1A-mutant cancers. Nat. Med. Nature Publishing Group, a division of Macmillan Publishers Limited. All Rights Reserved.; 2014;20:251–4.

28. Chandler RL, Brennan J, Schisler JC, Serber D, Patterson C, Magnuson T. ARID1a-DNA interactions are required for promoter occupancy by SWI/SNF. Mol. Cell. Biol. 2013;33:265–80.

29. Zhu P, Wang Y, Wu J, Huang G, Liu B, Ye B, et al. LncBRM initiates YAP1 signalling activation to drive self-renewal of liver cancer stem cells. Nat. Commun. 2016;7:13608.

30. Wang Y, He L, Du Y, Zhu P, Huang G, Luo J, et al. The long noncoding RNA lncTCF7 promotes self-renewal of human liver cancer stem cells through activation of Wnt signaling. Cell Stem Cell. 2015;16:413–25.

31. Marathe HG, Watkins-Chow DE, Weider M, Hoffmann A, Mehta G, Trivedi A, et al. BRG1 interacts with SOX10 to establish the melanocyte lineage and to promote differentiation. Nucleic Acids Res. [Internet]. 2017; Available from: http://dx.doi.org/10.1093/nar/gkx259

32. Park J, Wood MA, Cole MD. BAF53 forms distinct nuclear complexes and functions as a critical c-Myc-interacting nuclear cofactor for oncogenic transformation. Mol. Cell. Biol. 2002;22:1307–16.

33. Wang L, Zhao Z, Meyer MB, Saha S, Yu M, Guo A, et al. CARM1 methylates chromatin remodeling factor BAF155 to enhance tumor progression and metastasis. Cancer Cell. 2014;25:21–36.

34. Langmead B, Salzberg SL. Fast gapped-read alignment with Bowtie 2. Nat. Methods. 2012;9:357–9.

35. Li H, Handsaker B, Wysoker A, Fennell T, Ruan J, Homer N, et al. The Sequence Alignment/Map format and SAMtools. Bioinformatics. 2009;25:2078–9.

36. Ramírez F, Ryan DP, Grüning B, Bhardwaj V, Kilpert F, Richter AS, et al. deepTools2: a next generation web server for deep-sequencing data analysis. Nucleic Acids Res. 2016;44:W160–5.

37. Robinson JT, Thorvaldsdóttir H, Winckler W, Guttman M, Lander ES, Getz G, et al. Integrative genomics viewer. Nat. Biotechnol. 2011;29:24–6.

38. Zhang Y, Liu T, Meyer CA, Eeckhoute J, Johnson DS, Bernstein BE, et al. Model-based analysis of ChIP-Seq (MACS). Genome Biol. 2008;9:R137.

39. Bray NL, Pimentel H, Melsted P, Pachter L. Erratum: Near-optimal probabilistic RNA-seq quantification. Nat. Biotechnol. 2016;34:888.

40. Soneson C, Love MI, Robinson MD. Differential analyses for RNA-seq: transcript-level estimates improve gene-level inferences. F1000Res. 2015;4:1521.

41. Love M, Anders S, Huber W. Differential analysis of count data‐‐the DESeq2 package. Genome Biol. 2014;15:550.

42. Dykhuizen EC, Hargreaves DC, Miller EL, Cui K, Korshunov A, Kool M, et al. BAF complexes facilitate decatenation of DNA by topoisomerase IIα. Nature. 2013;497:624–7.

